# Modular evolution and functional divergence of the *Pseudomonas* Fit toxin locus

**DOI:** 10.64898/2026.11.17.763165

**Authors:** Jordan Vacheron, Kin Ki Jim, Géraldine Menoud, Clara M. Heiman, Daniel Garrido-Sanz, Jan-Willem Veening, Christoph Keel

## Abstract

The insecticidal Fit toxin is well known for its role in insect interactions in plant-associated *Pseudomonas*. However, its presence in non-entomopathogenic *Pseudomonas*, including the marine fish pathogen *Pseudomonas baetica*, raises questions about its evolutionary history and functional diversification. Here, we investigate the *fit* locus across the *Pseudomonas fluorescens* species complex using comparative genomic, structural, and functional approaches. Phylogenomic analyses support several ancient acquisitions of the *fit* gene cluster, followed by *Pseudomonas* subgroup-specific chromosomal integration and diversification of the toxin FitD and the sensor kinase FitF. Although *P. baetica* carries a *fit* operon, it showed no oral toxicity toward insect larvae. This lack of toxicity is associated with resistance to gut colonization by the commensal *Enterobacter* sp. Direct hemolymph injection also failed to reveal toxicity and the Fit activity in *P. baetica* is limited by the absence of *fit* promoter activation in insect hemolymph. Finally, in a zebrafish model, FitD is dispensable for *P. baetica*, which causes severe systemic infection through other virulence mechanisms. Nonetheless, FitD contributes to the virulence of *P. protegens* CHA0 in zebrafish, thereby extending its effects from insects to vertebrate hosts. Together, these findings highlight the evolutionary and regulatory plasticity of the *fit* locus and show how an ancestral toxin system can acquire different functional roles in bacteria adapted to distinct ecological niches.

## Introduction

Some bacteria belonging to the *Pseudomonas fluorescens* species complex are widespread inhabitants of soil and plant root environments. Many of these root-colonizing pseudomonads promote plant growth and suppress soil-borne fungal diseases through the production of diverse antimicrobial secondary metabolites, such as 2,4-diacetylphloroglucinol (DAPG), pyoluteorin, phenazines, and pyrrolnitrin^1–3^. Beyond their antifungal capabilities, specific biocontrol strains belonging to the *Pseudomonas protegens* and *Pseudomonas chlororaphis* subgroups exhibit potent insecticidal activity against agriculturally relevant pest insects, including lepidopteran larvae such as *Pieris brassicae*, *Spodoptera littoralis*, *Galleria mellonella*, and *Manduca sexta* ^4–6^.

This insecticidal phenotype carrid out by these strains is primarily mediated by a conserved 8-gene genomic cassette termed the *fit* locus (*fitABCDEFGH*, for *P. fluorescens* insecticidal toxin)^5^. The main product of this locus is FitD, a high-molecular-weight proteinaceous toxin (>300 kDa) that shares 73% amino acid sequence identity with the Makes caterpillars floppy 1 (Mcf1) toxin from *Photorhabdus luminescens*, a bacterial endosymbiont of insect-pathogenic entomophagous nematodes^5,7^. In *P. luminescens*, Mcf toxins act as primary virulence factors. Upon injection or release into the insect hemocoel, Mcf1 targets host phagocytic hemocytes and midgut epithelial cells, triggering rapid membrane blebbing and cell apoptosis via a pro-apoptotic BH3-like motif, an autoproteolytic cysteine protease domain, and a TcdA/TcdB-like translocation/pore-forming module ^8–11^. This causes massive tissue damage that results in a characteristic collapse of larval body turgor and in the distinctive “floppy” phenotype, melanization, and rapid death within 12–24 h^4–6^. In *P. protegens* CHA0, the FitD toxin is secreted by a dedicated Type I secretion system (FitABC and FitE) and tightly regulated by a three-component regulatory system consisting of the hybrid histidine kinase sensor FitF, the LysR-type transcriptional activator FitG, and the repressor FitH^5^. FitF senses host-derived chemical signals in insect hemolymph through its periplasmic double-PASp domain, driving strong transcriptional activation from the Pfit promoter specifically inside the host^4,12^.

For nearly two decades since its initial discovery, extensive genomic screens and phylogenetic analyses had established that the *fit* gene cluster was exclusively restricted to plant-colonizing pseudomonads belonging to the *P. protegens* and *P. chlororaphis* subgroups^6^. Consequently, the prevailing paradigm held that the Fit toxin represented a highly specialized evolutionary adaptation confined to terrestrial, root-associated bacteria, enabling them to exploit soil-dwelling insects as an alternative host niche or defend their rhizosphere habitat against root-feeding herbivores^13–15^.

In this study, we challenge this paradigm by systematically investigating the broader genomic distribution, evolutionary history, and functional significance of the insecticidal Fit toxin cluster across the *Pseudomonas* genus. Our phylogenomic analysis revealed the presence of a complete, structurally conserved *fitABCDEFGH* operon in free-living soil isolates (*Pseudomonas* sp. Irchel s3a18 and s3b6) that are phylogenetically distinct from the *P. protegens* and *P. chlororaphis* subgroups. The operon was also found in *Pseudomonas baetica* LMG25716, a psychrophilic marine fish pathogen isolated from an environment unrelated to plant roots that causes systemic disease in farmed flatfish (wedge sole, *Dicologlossa cuneata*)^16,17^. To understand the biological significance and evolutionary implications of harboring an insecticidal toxin cluster in a fish pathogen, we combined phylogenomic analyses, comparisons of protein domain architecture, transcriptional assays, and *in vivo* infection experiments in the pest insect *Pieris brassicae* as well as zebrafish (*Danio rerio*) embryos. We demonstrated that despite the presence of the *fit* gene cluster, *P. baetica* and Irchel strains lack toxicity toward insects due to microbiota-mediated gut exclusion by commensal *Enterobacter* sp. and sensory rewiring of the FitF sensor kinase. Importantly, we show for the first time that *P. protegens* CHA0 is virulent in a zebrafish embryo infection model and that FitD contributes to this phenotype, extending the known role of this toxin beyond insect hosts. By contrast, *P. baetica* showed greater systemic killing activity than CHA0 after direct vascular administration, but survival did not differ significantly between embryos infected with wild-type *P. baetica* and those infected with its Δ*fitD* mutant, suggesting that other virulence factors may drive pathogenicity in this species.

## Results and discussion

### The insecticidal Fit toxin gene cluster is widely distributed across diverse *Pseudomonas* lineages beyond plant-beneficial biocontrol strains

To determine how the *fit* gene cluster is distributed across the genus, we mapped its presence into the *Pseudomonas* species complex (**Figure 1**). The cluster was detected in only four of the eleven subgroups represented: the *P. protegens*, *P. chlororaphis*, *P. koreensis* and a in new subgroup represented by two bacterial strains, designated in this study as the “Irchel” subgroup. Even within the four positive subgroups, the distribution was discontinuous. The distribution of the *fit* cluster varied within subgroups previously known to contain Fit-producing strains (**Figure 1**). Within the *P. chlororaphis* subgroup, while the *fit* cluster is conserved in *P. chlororaphis* subsp. *aureofaciens* and *P. chlororaphis* subsp. *piscium*, it is absent in strains belonging to *P. chlororaphis* subsp. *aurantiaca* and *P. chlororaphis* subsp. *chlororaphis*. Similarly, occasional losses of the *fit* gene cluster occurred within the *P. protegens* subgroup (e.g., *P. saponiphila* DSM 9751 or LD120), aligned with previous observations of evolutionary plasticity and local cluster dynamics within this subgroup^18^.

**Figure 1:**
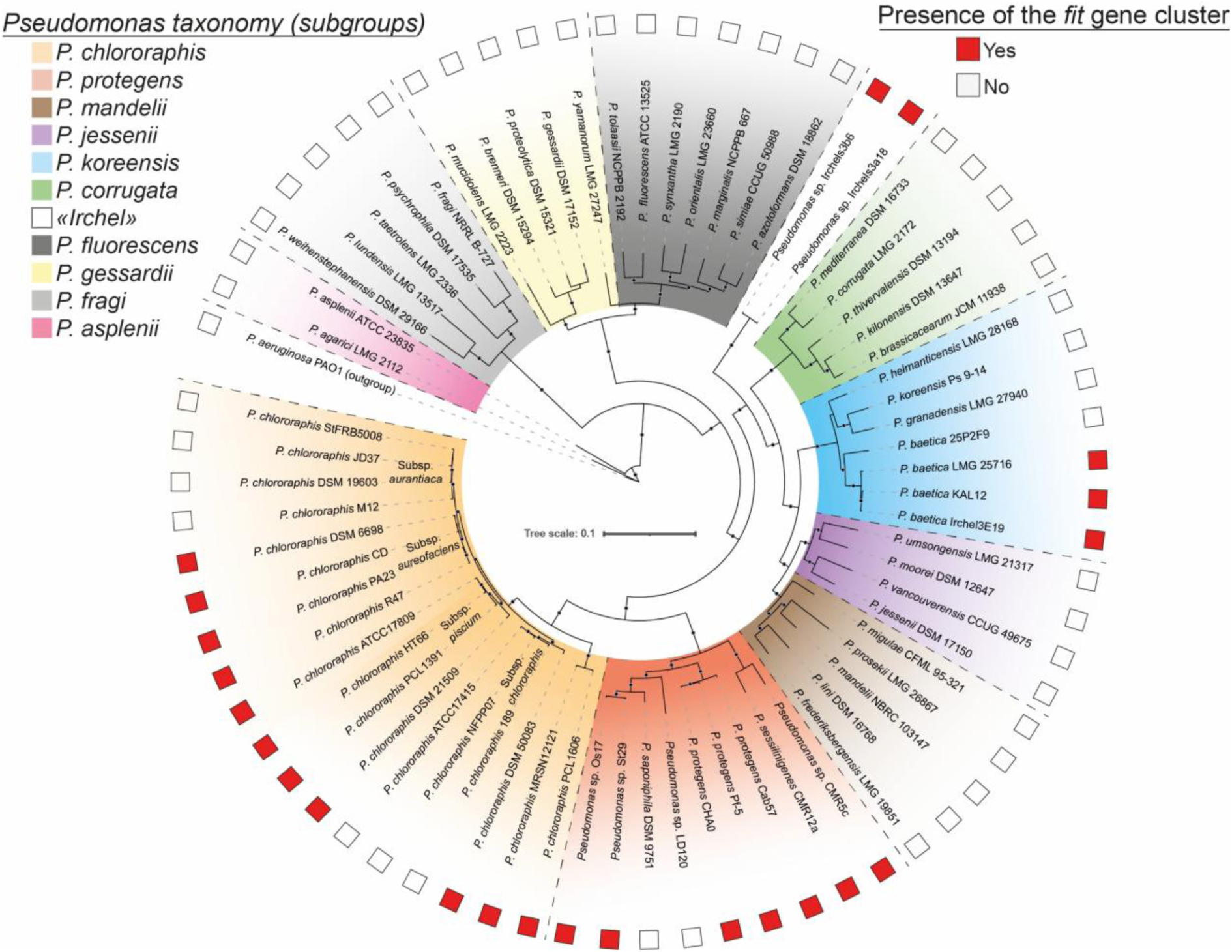
Distribution of the *fit* gene cluster across diverse subgroups of the *Pseudomonas* species complex. Circular maximum-likelihood phylogenetic tree inferred from aligned concatenated core protein homologs shared across representative genomes of the *Pseudomonas fluorescens* species complex, rooted using *Pseudomonas aeruginosa* PAO1 as the outgroup. Shaded background sectors delineate recognized *Pseudomonas* subgroups. The unshaded sector highlights the newly defined Irchel subgroup. Red squares on the outer circumference indicate the presence of the complete 8 genes *fitABCDEFGH* cluster; open squares denote its absence.

Furthermore, the *fit* cluster was uncovered in environmental soil isolates from Switzerland, specifically *Pseudomonas* sp. Irchel s3a18 and s3b6^19^. These strains form a coherent, distinct, and uncharacterized clade within the *P. fluorescens* species complex, designated here as the “Irchel” subgroup (**Figure 1**). The full *fit* operon was also identified in the genomes of several members of the bacterial species *P. baetica*, belonging to the *P. koreensis* subgroup, including the type strain LMG25716. This type strain is a temperature-dependent fish pathogen originally isolated from the liver of diseased wedge sole (*D. cuneata*) in marine aquaculture, where it causes systemic pathology and high mortality^17^.

The presence of a conserved *fitABCDEFGH* locus across such ecologically heterogeneous bacteria ranging from plant-beneficial root-colonizers (*P. protegens and P. chlororaphis*) and environmental soil isolates (*Irchel* group) to aquatic fish pathogens (*P. baetica*) challenges the established paradigm that the Fit toxin function is intrinsically linked to plant-protective or purely entomopathogenic lifestyles^6,13^. This gene distribution raises questions regarding the evolutionary origins of the locus, its genomic context, and its functional role or potential host specificity across these diverse ecological niches.

### The *fit* locus is structurally conserved but occupies subgroup-specific genomic regions

To determine the genomic organization and structural integrity of the *fit* locus, we examined its genomic neighborhood across the 17 representative *Pseudomonas* genomes (**Figure 2**). In all genomes, the gene cluster spans approximately 22 kb and comprises the exact same eight genes (*fitA*–*fitH*) in an identical order and orientation (**Figure 2**). Thus, the *fit* cluster itself constitutes a well-defined, structurally conserved genetic unit wherever it occurs within the *Pseudomonas* genus.

**Figure 2:**
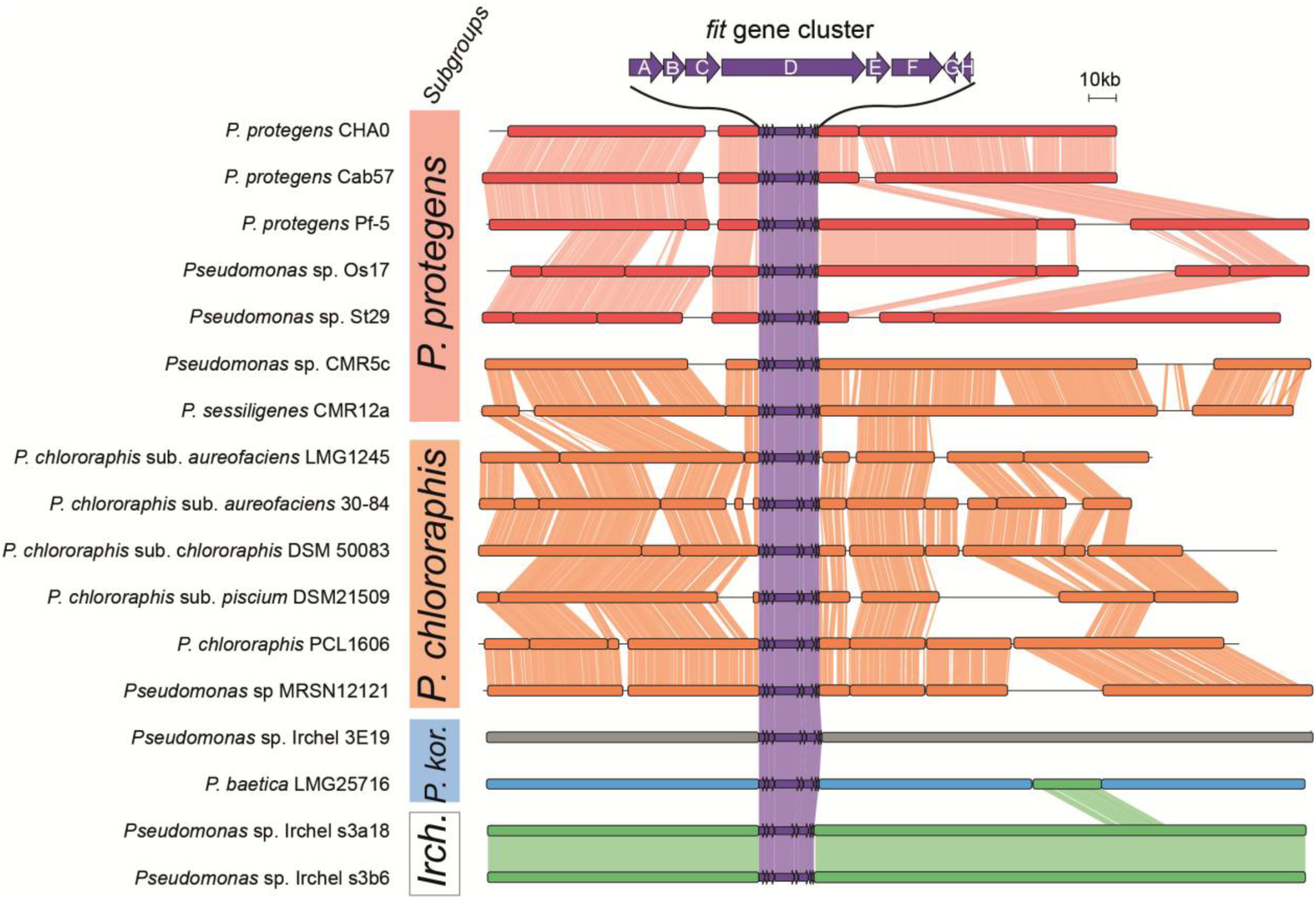
Genomic synteny and conserved locus architecture of the *fit* gene cluster across *Pseudomonas* subgroups. Alignment of the *fit* gene cluster (*fitA–fitH*, shown in purple) and surrounding genomic regions across representative genomes of the *Pseudomonas fluorescens* species complex. Gene representation is simplified by grouping genes into blocks based on their similarity to those in adjacent genomes; full representation is available in **supplementary data 1**. Background shading delineates taxonomic subgroups. Connecting ribbons indicate regions of high sequence conservation and conserved synteny (>70% amino acid identity) between adjacent genomes. *P. kor*: *P. koreensis* ; *Irch*: Irchel.

In contrast to the highly conserved internal organization of the *fit* gene cluster, the chromosomal regions flanking the locus were conserved strictly within individual taxonomic subgroups (**Figure 2**). In the *P. chlororaphis* subgroup, all six analyzed strains displayed complete genomic synteny on both sides of the cluster. In the *P. protegens* subgroup, the flanking regions were likewise syntenic among strains CHA0, cab57, Pf-5, Os17, and St29. However, the region located immediately downstream of the cluster exhibited local structural variability: strains Pf-5 and Os17 shared one distinct arrangement corresponding to the presence of a cluster of gene encoding for the rhizotoxin^18^ (**Figure 2**). The two strains from the *Irchel* subgroup were fully colinear over the entire chromosomal region analyzed. In the *P. koreensis* subgroup, the genes flanking the cluster in strains 3E19 and LMG25716 showed no detectable sequence homology with each other, nor with those of the other subgroups, with the sole exception of a short segment downstream of the cluster in *P. baetica* LMG25716 that is homologous to a genomic region in the genomes of the Irchel strains (**Figure 2**). Beyond these subgroup-specific arrangements, no overarching synteny was detected between the flanking regions of distinct subgroups, indicating that each subgroup carries the *fit* cluster at a different chromosomal insertion site.

The single exception concerns *Pseudomonas* sp. CMR5c and *Pseudomonas sessilinigenes* CMR12a. Although both strains unequivocally belong to the *P. protegens* subgroup according to the species-level phylogenetic tree (**Figure 1**), their *fit* cluster is detected within a *chlororaphis*-type genomic context (**Figure 2**). Specifically, the flanking regions of CMR5c and CMR12a are syntenic with those of *P. chlororaphis* strains over approximately 100 kb on both sides of the cluster, rather than displaying the *protegens*-type flanking backbone.

The evolutionary divergence of the cluster-encoded proteins closely mirrored these genomic patterns (**Figure 2**). Average amino-acid identities of the *fit* gene cluster-encoded proteins, as determined by clinker analysis, ranged from 86% to 88% among closely related strains within a given subgroup, decreased to 73-79% between the *P. protegens* and *P. chlororaphis* subgroups, and dropped to 64-69% when comparing these two subgroups to the *P. koreensis* or Irchel subgroups. However, the cluster-encoded proteins of the CMR5c and CMR12a strains shared a slightly higher amino-acid identity with *P. chlororaphis* strains (average of 79%) than with other *P. protegens* strains (average of 74%), a pattern consistent with a possible ancestral horizontal locus replacement or recombination event in these two bacterial strains.

Together, these results show that the *fit* cluster is internally conserved but embedded in distinct, subgroup-specific genomic contexts, with CMR5c and CMR12a displaying an atypical *P. chlororaphis*-like configuration. This evolutionary and genomic heterogeneity prompted us to examine whether the toxin itself also exhibits subgroup-specific variation, focusing on the domain architecture of the toxin FitD.

### The Fit toxin cluster of *Pseudomonas* derives from at least three acquisitions from *Photorhabdus*/*Xenorhabdus*-related donors

The toxin FitD was previously described to be homologous to the Mcf (“makes caterpillars floppy”) toxins of the insect pathogens *Photorhabdus* and *Xenorhabdus*^5,20^. When focusing on the domain architecture of FitD, it differed markedly between subgroups (**Figure 3**). In *P. protegens* and *P. chlororaphis* strains, FitD (2993-3004 aa) had the same architecture as Mcf1 of *Photorhabdus laumondii* TTO1 (2997 aa), with an N-terminal Mcf1 effector domain (PF30645) followed by the activator-binding (PF30646), protease effector (PF30660) and TcdA/TcdB pore-forming (PF12920) domains.

**Figure 3:**
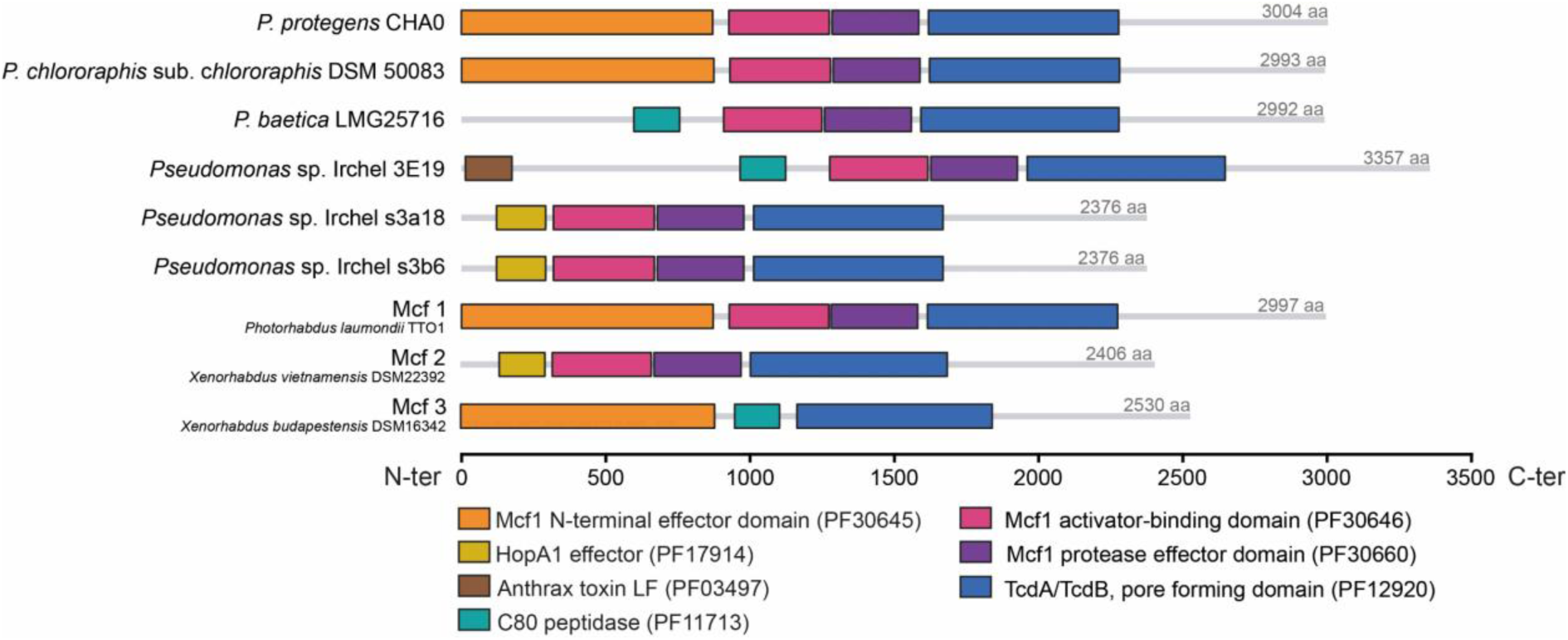
Modular domain architecture and structural variation of FitD toxin variants and related Mcf proteins. Schematic representation of the protein domain architectures of FitD toxin variants from representative *Pseudomonas* from *the P. protegens* subgroup (CHA0)*, P. chlororaphis* subgroup (DSM50083)*, P. koreensis* subgroup (LMG25716 and Irchel 3E19) and the “Irchel” subgroup (Irchel s3b6 and Irchel s3a18). Protein lengths in amino acids (aa) are indicated on the right. Colored boxes correspond to functional Pfam domains predicted by InterpProScan^34^.

While plant-associated *Pseudomonas protegens* and *Pseudomonas chlororaphis* strains harbor the canonical Mcf1-like effector architecture^5^, the incorporation of distinct, highly specialized catalytic and regulatory modules in *Pseudomonas baetica* and the Irchel strains points to a potentially altered host interaction mechanisms and functional diversification.

In the soil-dwelling Irchel strains, FitD lacks the Mcf1 effector domain and instead carries an N-terminal HopA1-like domain (PF17914), mirroring the domain organization of Mcf2 in *Xenorhabdus vietnamensis* (**Figure 3** ^20^). HopA1, originally identified as HrmA in *Pseudomonas syringae*, is a well-characterized Type III secretion system effector in phytopathogenic bacteria that targets host signaling cascades, particularly mitogen-activated protein kinases, to suppress plant innate immunity^8^. The presence of this HopA1 domain may act as a translocated effector targeting eukaryotic kinase pathways or immune regulators in soil microeukaryotes or invertebrates, although this hypothesis remains to be tested experimentally.

FitD of *P. baetica* LMG25716 and of strain 3E19 also carries a peptidase C80 domain (PF11713), which is absent from the *P. protegens*, *P. chlororaphis* and Irchel variants but present in Mcf3 of *Xenorhabdus budapestensis* (**Figure 3**). This domain belongs to the cysteine protease domain (CPD) of MARTX toxins, which is activated by the eukaryote-specific inositol hexakisphosphate and cleaves the toxin to release its effectors^21^. A related Mcf-like effector of MARTX toxins is likewise activated by Arf proteins and cleaves Rab GTPases^22,23^. The C80 domain of the *P. baetica* FitD might therefore provide an additional, InsP6-dependent autoprocessing step, in addition to or instead of the Arf-dependent activation.

To determine whether these distinct FitD domain architectures reflect different evolutionary origins, we next examined the phylogenetic relationships among FitD and Mcf proteins (**Figure 4**). FitD of *P. protegens* and *P. chlororaphis* strains grouped with the Mcf1 clade of *Photorhabdus*, whereas those of the *P. baetica* and Irchel strains grouped with the Mcf2 clade, and Mcf3 formed a separate one. FitD of *Pseudomonas* is therefore polyphyletic, which suggests several independent acquisition events from donors related to *Photorhabdus* and *Xenorhabdus*. The *P. protegens* and *P. chlororaphis* sequences, which are closely related to each other, grouped with the Mcf1 clade, and were probably acquired once, from a source related to Mcf1. The Fit cluster of *P. baetica* strains from the *P. koreensis* subgroup and the Irchel strains from the Irchel subgroup both grouped with the Mcf2 clade, but these two subgroups are separated in the species phylogeny by other subgroups lacking the cluster (**Figure 1**), so that inheritance from a common ancestor would require multiple independent losses. A more parsimonious scenario is that the cluster was acquired independently in each of the two subgroups, from donors related to the Mcf2 clade. Moreover, the average amino-acid identity between the FitABCDEFGH proteins of the *P. protegens P. chlororaphis* subgroups and the one of the *P. baetica*/Irchel subgroup was low (64-69%), and the amino-acid identity between the FitABCDEFGH proteins of the *P. baetica* strains and those of the Irchel subgroup was similarly low (64-65%) which is consistent with the distinct origins of the Fit cluster in these subgroups.

**Figure 4.**
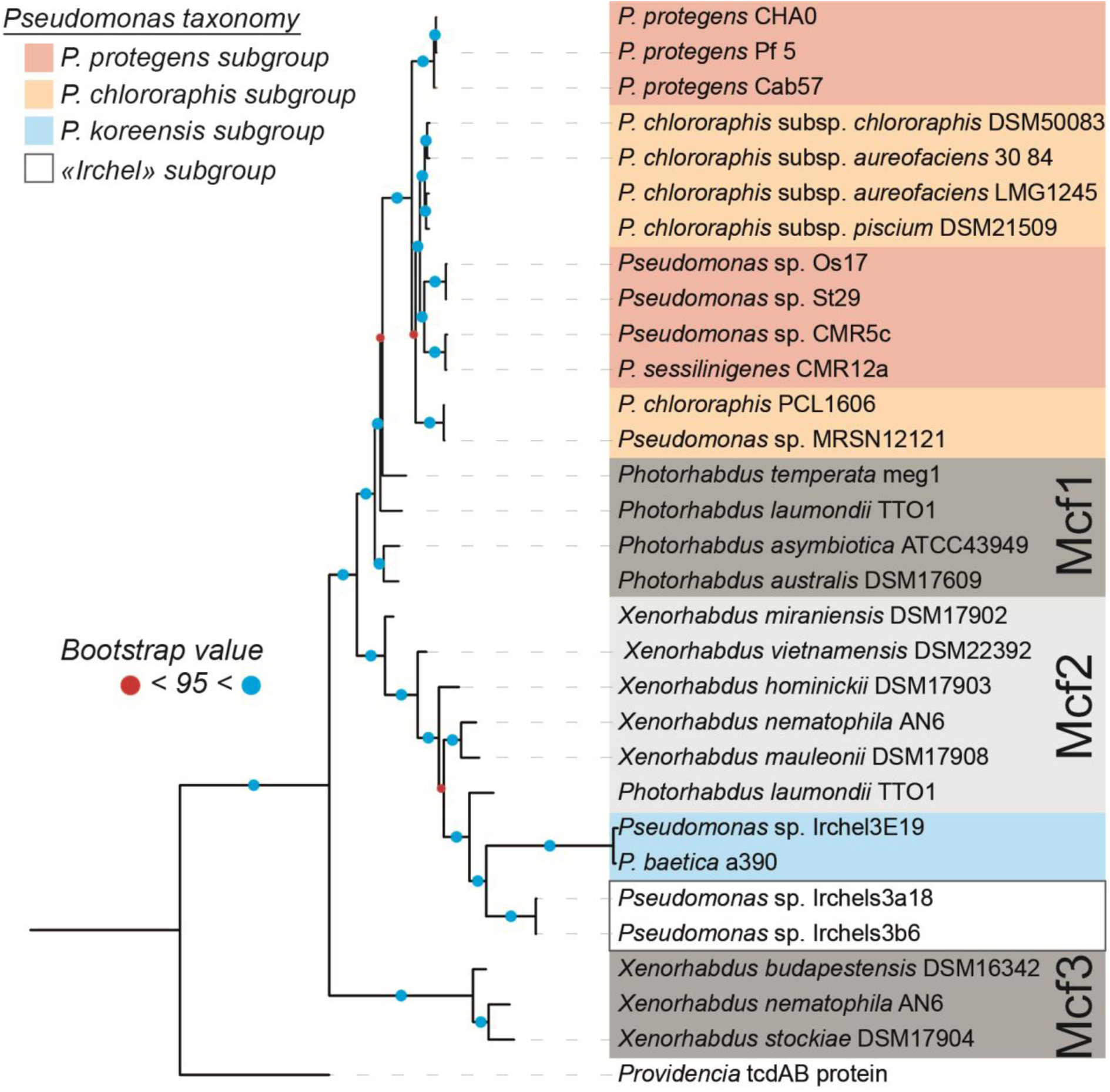
Phylogenetic tree of full length FitD/Mcf proteins. The tree was rooted using the TcdA/TcdB protein sequence of *Providencia* as an outgroup. Shaded background boxes correspond to different *Pseudomonas* subgroups and toxin clades (*mcf1*, *mcf2*, and *mcf3* clades according to Ruffner and colleagues^20^). Blue circles at nodes indicate bootstrap support values ≥ 95%; red circles indicate bootstrap values < 95%.

Variation in the N-terminal region partially followed these phylogenetic relationships (**Figure 3**). Although *P. protegens* and *P. chlororaphis* FitD share the Mcf1-type effector domain, and Irchel FitD carry a HopA1 domain, as in Mcf2, the phylogenetic pattern is less clear in with the *P. baetica* strains. Indeed, their FitD proteins group with the Mcf2-like lineage but they carry distinct N-terminal modules (**Figure 3**).

Beyond FitD, the evolution of the whole Fit cluster is not explained by vertical inheritance within the entire *Pseudomonas fluorescens* species complex. The cluster is present in only four subgroups and absent from closely related genomes within them (**Figure 1**), and it lies in a different genomic context in each subgroup (**Figure 2**). This suggests that, after acquisition, the cluster was inherited vertically within each subgroup / bacterial species, with sporadic losses explaining its patchy distribution. However, the phylogenies constructed with the other *fit* genes were discordant with the species phylogeny (**Supplementary Figure 1**), and *P. protegens* and *P. chlororaphis* sequences were interspersed. Because these two subgroups carry the cluster at different loci while sharing similar proteins (73-79% aa identity for the entire cluster), the cluster has probably also been exchanged between *Pseudomonas* from distinct subgroups. The clearest case of this hypothesis is with CMR5c and CMR12a. Although these two strains belong to the *P. protegens* subgroup^18^ (**Figure 1**), both the cluster and a flanking sequence of about 100 kb of resemble those of *P. chlororaphis* strains(**Figure 2**), a pattern that might result from homologous recombination of a large *P. chlororaphis*-derived fragment into a *P. protegens* recipient.

In summary, the Fit cluster has a patchy distribution in *Pseudomonas* strains, derives from at least three independent acquisitions (in the *P. protegens*/ *P. chlororaphis* ancestor, in *P. baetica* strains ancestor and in Irchel strains ancestor) from donors related to *Photorhabdus* and *Xenorhabdus*, and has been further shaped by gains, losses and potentially domain shuffling between *Pseudomonas* from different subgroups (*P. protegens* and *P. chlororaphis*). Among the resulting FitD variants, the one of *P. baetica* is particularly interesting. It groups with the Mcf2-like lineage but lacks the Mcf1 effector domain found in the FitD variant of *P. protegens* and *P. chlororaphis* and instead carries a C80 peptidase domain. We then tested whether *P. baetica* retains insecticidal activity despite the distinct domain architecture of its FitD toxin.

### *Pseudomonas baetica* lacks oral insecticidal toxicity due to microbiota-mediated gut colonization resistance

In insect-killing pseudomonads such as *P. protegens* CHA0, oral pathogenesis against herbivorous insect pests relies on a multifactorial virulence arsenal (^5,24–27^). After ingestion, CHA0 establishes itself in the insect digestive tract and uses its type VI secretion system (T6SS) to target and eliminate Gram-negative commensal bacteria, notably *Enterobacter* sp.^24^. The bacterium may then breach the gut epithelial barrier and enter the hemocoel through the action of virulence factors such as secreted lytic enzymes, cyclic lipopeptides, or ExlA-like two-partner secretion systems^25,26^. Once in the hemolymph, CHA0 evades insect immune defenses through specific O-antigen decorations (OSA) that confer resistance to antimicrobial peptides^12^. Host-derived signals are perceived by the hybrid histidine kinase FitF, activating the FitH/FitG regulatory cascade, strongly derepressing the *fitABCDE* promoter, and promoting the secretion of the proteinaceous toxin FitD^4,12^. FitD causes severe cytotoxicity, loss of body turgor (the “floppy” phenotype) and lethal septicemia, ultimately resulting in rapid insect death^5,7^.

Because the marine fish pathogen *P. baetica* LMG25716 harbors a structurally complete *fitABCDEFGH* gene cluster with the same gene order as the one belonging to *P. protegens* CHA0, we investigated whether it retains the ability to infect and kill insect pests. We therefore performed oral feeding assays with second-instar larvae of the cabbage butterfly *Pieris brassicae*, using *P. protegens* CHA0 as a positive control (**Figure 5**).

**Figure 5.**
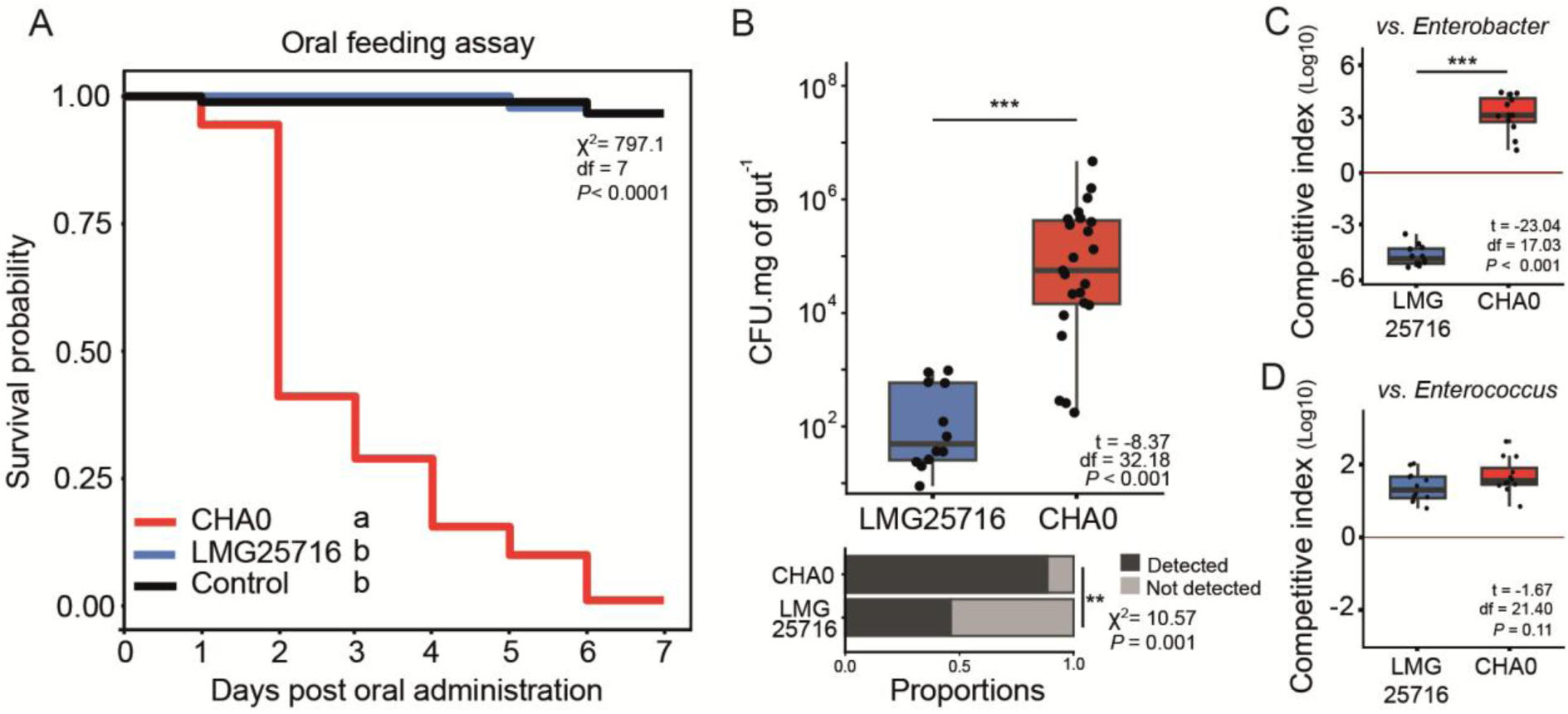
*Pseudomonas baetica* LMG 25716 lacks oral insecticidal activity against *Pieris brassicae* due to gut colonization failure and competitive exclusion by commensal *Enterobacter* sp. **(A)** Kaplan-Meier survival curves of *Pieris brassicae* 2nd-instar larvae fed on artificial diet inoculated with the different *Pseudomonas strains* (10^6^ cells) or sterile NaCl as control. Larvae survival was monitored daily for 1 week. Statistical significance was evaluated using a mixed-effect Cox proportional hazards model and different letters denote statistically significant differences at *P* < 0.05). **(B)** Quantification of bacterial colonization in individual *P. brassicae* larvae 24 h post oral administration. Box plots depict median bacterial loads (dots represent individual larvae). Bottom stacked bar plot displays the proportion of larvae in which inoculant bacteria were detected (dark grey) vs. not detected (light grey). Statistical significance was assessed using Welch’s two-sample *t*-test for colonization data and using Fisher’s exact test to compare proportion of colonized insects. (C, D) *In vitro* pairwise competition assays of *P. baetica* LMG 25716 and *P. protegens* CHA0 against gut commensal isolates *Enterobacter* sp. **(C)** and *Enterococcus* sp. **(D)** isolated from *P. brassicae* midguts. The horizontal red line represents competitive parity. Box plots represent data from three independent experiments with four technical replicates each. Statistical significance was assessed using Welch’s two-sample *t*-test.

Oral ingestion of *P. protegens* CHA0 caused rapid and lethal disease, resulting in 100% larval mortality within six days (**Figure 5A**). In contrast, *P. baetica* LMG 25716 showed no detectable oral toxicity toward *P. brassicae* (**Figure 5A**). This non-lethal phenotype was shared by the environmental soil isolates of the Irchel subgroup (**Supplementary Figure 2**).

To determine whether the absence of oral toxicity reflected a failure of *P. baetica* to establish within the host, we quantified bacterial loads in the larval midgut 24 h after infection (**Figure 5B**). *P. protegens* CHA0 established dense intestinal populations, with a median load of approximately 10^5^ CFU mg^−1^ of gut and detection in more than 90% of the larvae. By contrast, *P. baetica* LMG25716 remained near or below the detection limit (<10^2^ CFU mg^−1^ of gut) and was completely cleared from more than 50% of the larvae (**Figure 5B**). We hypothesized that the inability of *P. baetica* to colonize the insect digestive tract resulted from competitive exclusion by the indigenous gut microbiota of *P. brassicae*, which is dominated by two commensal taxa, *Enterobacter* sp. and *Enterococcus* sp.^24^. To test this hypothesis, we performed pairwise *in vitro* competition assays against each commensal (**Figure 5C, D**).

When competed against *Enterobacter* sp., *P. baetica* LMG25716 exhibited a strong competitive disadvantage, with a competitive index of approximately log_10_ CI = −5 (**Figure 5C**). By contrast, *P. protegens* CHA0 strongly suppressed *Enterobacter* sp., reaching approximately log_10_ CI = +3.5, as previously described^24^. In competition with the Gram-positive commensal *Enterococcus* sp., both *P. baetica* LMG25716 and *P. protegens* CHA0 maintained positive competitive indices of approximately log_10_ CI = +1.5 (**Figure 5D**). No significant difference was detected between the two strains indicating that their competitive performance against *Enterococcus* sp. was comparable.

Together, these results indicate that *P. baetica* encounters a microbiota-mediated colonization barrier in the lepidopteran gut that is specifically associated with commensal *Enterobacter* sp. Whereas entomopathogenic *P. protegens* CHA0 can suppress this competitor, *P. baetica* lacks the competitive capacity required to displace resident bacteria. This competitive disadvantage likely prevents initial gut establishment and, consequently, subsequent entry into the hemocoel.

### Bypassing the gut barrier does not reveal systemic toxicity in *Pseudomonas baetica* and fails to activate the *fit* promoter

Because *P. baetica* encounters a microbiota-mediated colonization barrier in the lepidopteran digestive tract (**Figure 5**), we next asked whether bypassing this barrier would reveal systemic entomopathogenicity. We therefore tested whether *P. baetica* could kill larvae after direct entry into the hemocoel or whether it remained intrinsically harmless to the insect host (**Figure 6**).

**Figure 6.**
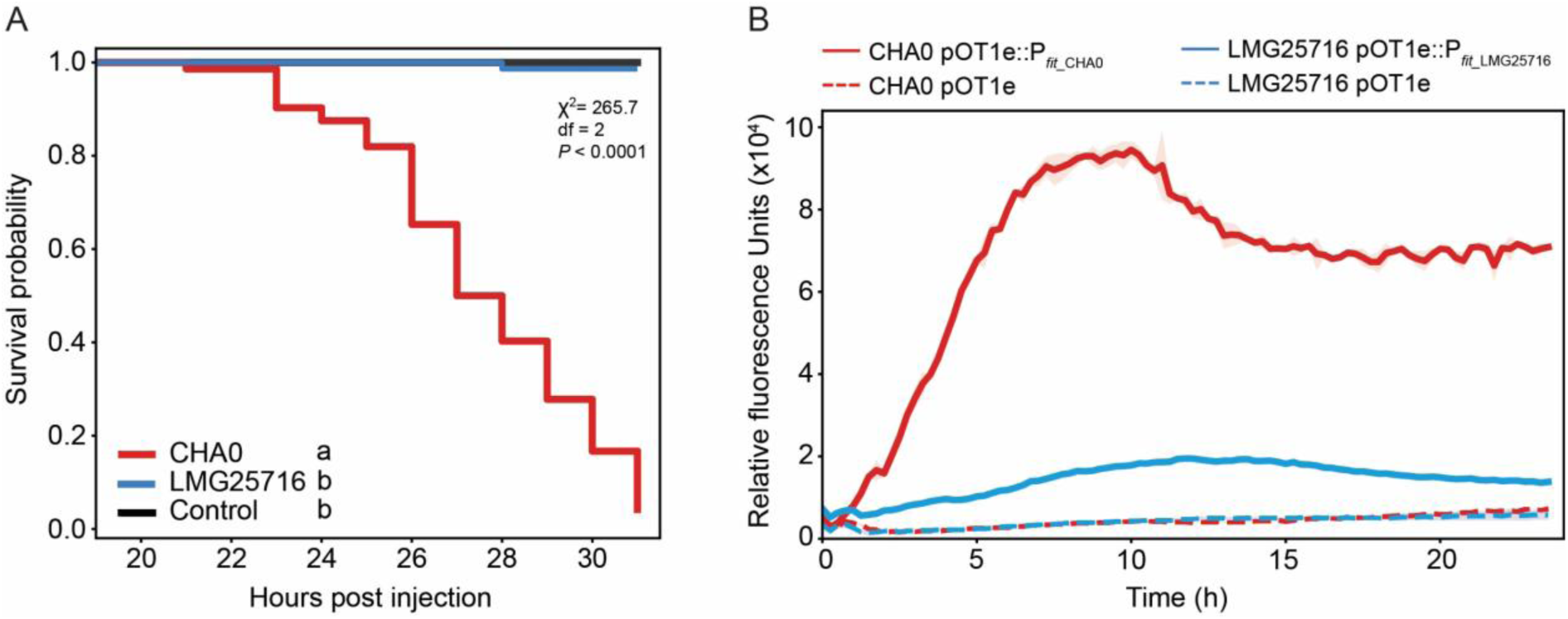
Bypassing the insect gut barrier confirms the absence of systemic infection of *Pseudomonas baetica* LMG 25716 due to non-activation of the *fitABCDE* promoter in insect hemolymph. **(A)** Direct injection of 10^2^ CFU of *Pseudomonas protegens* CHA0 and *Pseudomonas baetica* LMG 25716, 0.9% NaCl solution as control into the hemolymph of ultimate-instar *Pieris brassicae* larvae (n = 18 larvae per treatment). Kaplan-Meier survival curves depict the probability of larval survival over a 32 h post-injection timeline. Statistical significance was determined using a mixed-effect Cox proportional hazards model; asterisks *** indicate *P* < 0.001. **(B)** Real-time kinetic monitoring of P*_fitABCDE_* promoter activity (noted P*_fitA_*) in *P. protegens* CHA0 and *P. baetica* LMG 25716 during growth in Grace’s Insect Medium (GIM) over 24 h. Strains harboring the promoterless pOT1e vector served as negative controls and represented with dashed lines. Data are represented as the mean ± standard deviation of relative fluorescence units normalized by cell density (OD_600nm_) from three independent biological replicates with three technical replicates each.

To assess systemic toxicity, we injected approximately 10^2^ CFU per larva of *P. baetica* LMG25716 or *P. protegens* CHA0 directly into the hemocoel of ultimate-instar *P. brassicae* larvae (**Figure 6A**). As expected, injection of *P. protegens* CHA0 was rapidly lethal, causing 100% mortality within 30 h post injection. By contrast, *P. baetica* LMG25716 did not cause mortality during the monitoring period (**Figure 6A**). We then examined whether this absence of systemic toxicity was associated with a failure to activate the *fitABCDE* promoter. Promoter activity was monitored using GFP-based transcriptional reporters during growth in Grace’s Insect Medium (GIM), a defined medium rich in amino acids and C4-dicarboxylates that mimics lepidopteran hemolymph (**Figure 6B**). In *P. protegens* CHA0, growth in GIM strongly induced the *fitABCDE* promoter, producing a greater than 14-fold increase in relative fluorescence that peaked during early stationary phase. By contrast, promoter activity in *P. baetica* LMG 25716 remained at basal levels throughout the 24 h of incubation, with no detectable induction (**Figure 6B**).

Domain analysis showed that FitF of CHA0 (1072 aa) and *P. baetica* LMG25716 (1225 aa) share the architecture of a hybrid histidine kinase, with two transmembrane helices flanking a periplasmic Cache/PDC-type sensor domain (about 274 aa in both), followed by a cytoplasmic PAS domain and HisKA, HATPase, receiver and Hpt modules (**Figure 7**). The main difference is a putative second PAS-fold domain in the cytoplasmic region of *P. baetica* FitF (residues 509-592), which is absent from CHA0 FitF. The periplasmic sensor domains share 55% amino-acid identity, suggesting that the signals perceived by FitF may differ between the two strains.

**Figure 7.**
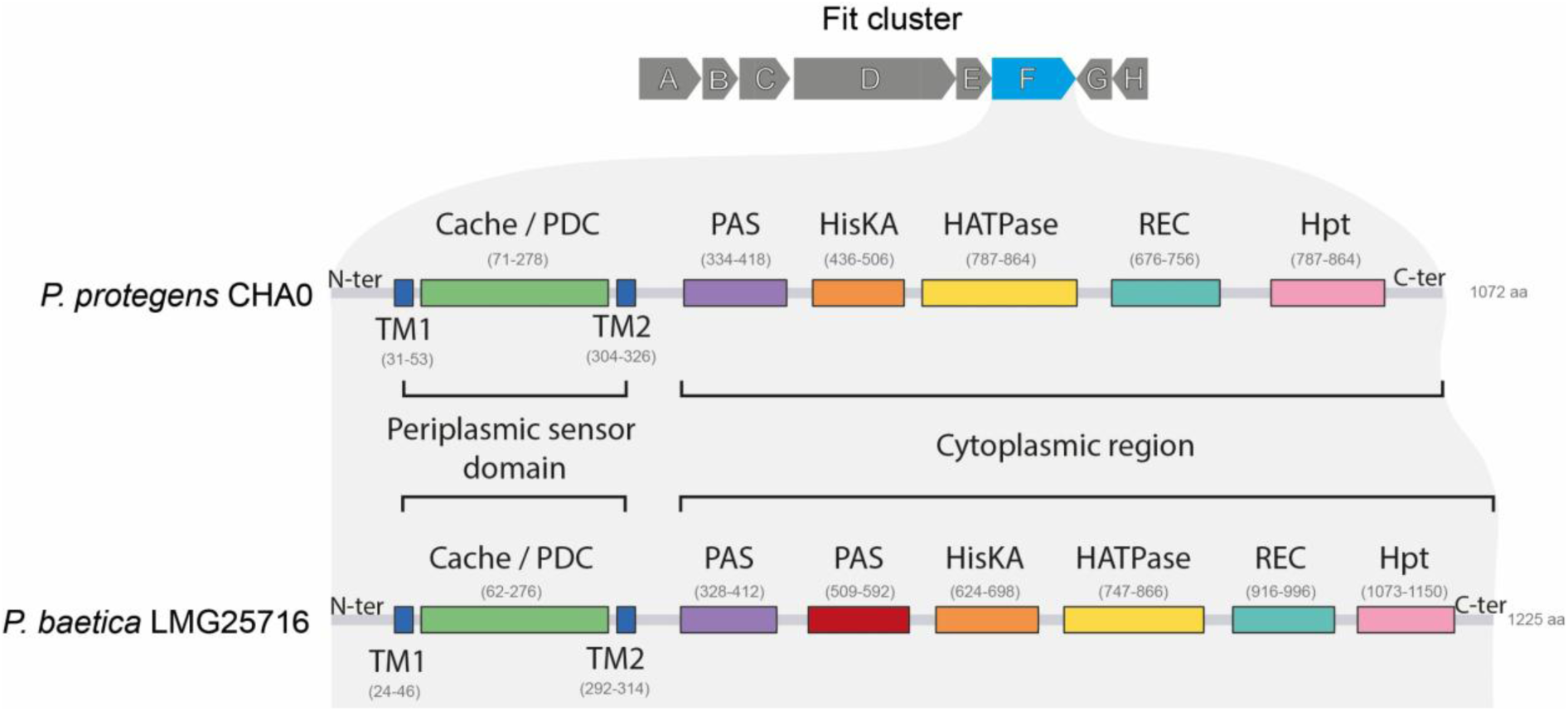
Domain architecture and structural organization of the FitF sensor protein in *Pseudomonas protegens* CHA0 and *Pseudomonas baetica* LMG25716. Schematic comparison of the protein domain architectures of the hybrid histidine kinase sensor FitF (encoded within the 8-gene *fitABCDEFGH* cluster, top) from *Pseudomonas protegens* CHA0 and *Pseudomonas baetica* LMG25716. TM, transmembrane helix; Cache, calcium channels and chemotaxis receptors; PDC, PhoQ-DcuS-CitA; PAS, Per-Arnt-Sim; HisKA, histidine kinase A; HATPase, histidine kinase-like ATPase; REC, receiver domain; Hpt, histidine-containing phosphotransfer domain.

Together, these findings show that the absence of insecticidal activity in *P. baetica* cannot be explained solely by its clearance from the midgut. Even after direct delivery into the hemocoel, the bacterium remained nonlethal, and its *fitABCDE* promoter was not induced under conditions that mimic insect hemolymph. Combined with the sequence divergence observed in the sensory domain of FitF, these results suggest that regulation of the Fit toxin may be adapted to signals associated with a different host or environment. Because *P. baetica* LMG25716 is a marine fish pathogen that causes systemic mortality in teleost species^16,17^, the *fitABCDEFGH* cluster may function in aquatic or vertebrate host contexts rather than in the insect hemocoel. Determining the contribution of FitD to interactions with fish and to aquatic host pathogenicity will therefore be important for defining the biological role of this toxin cluster in *P. baetica*.

### FitD contributes to zebrafish embryo mortality in *Pseudomonas protegens* but is dispensable for *Pseudomonas baetica* virulence

To determine whether FitD contributes to vertebrate pathogenesis and whether its role differs between plant-associated and aquatic lineages, we compared *P. protegens* CHA0, *P. baetica* LMG25716, and their respective isogenic Δ*fitD* mutants in zebrafish (*D. rerio*) embryos. Virulence was assessed using both bath immersion and systemic infection by caudal vein injection (**Figure 8**).

**Figure 8.**
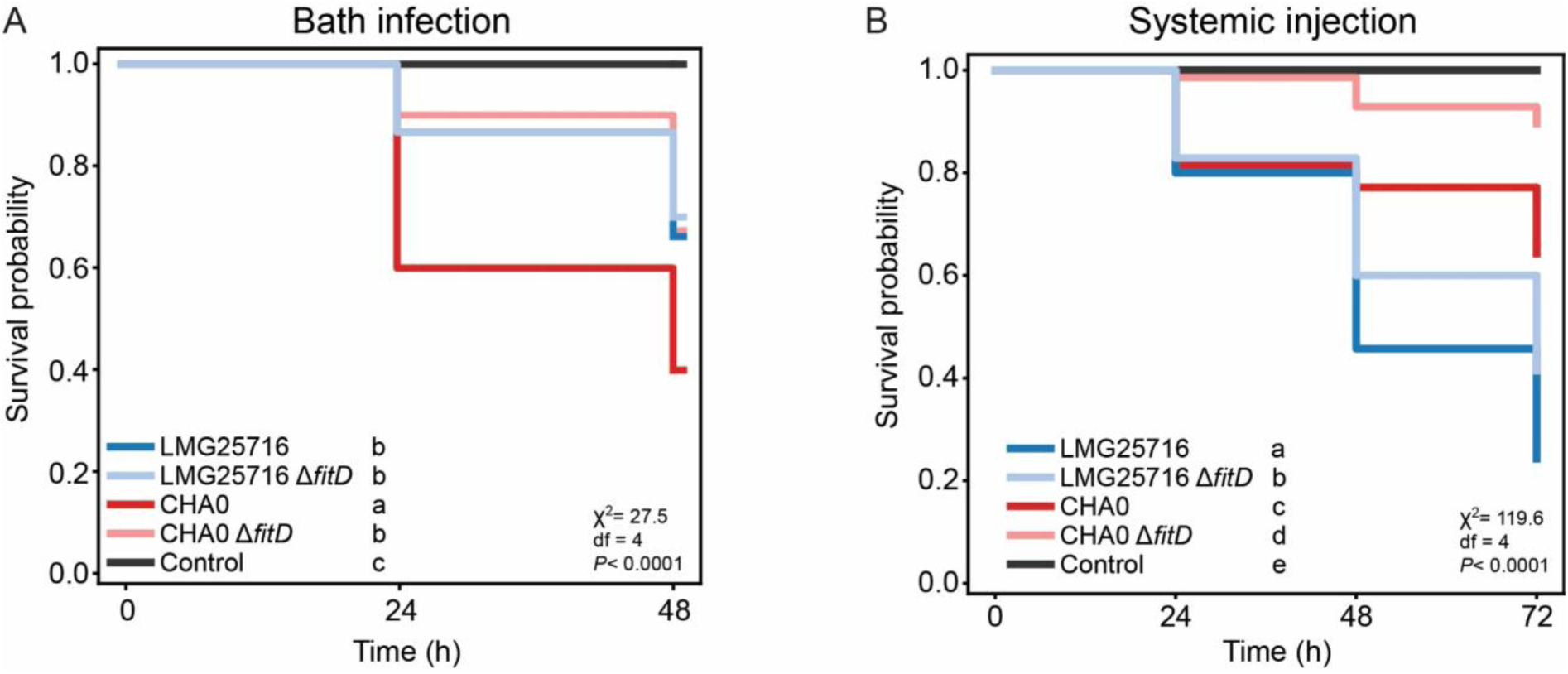
Divergent contribution of the FitD toxin to zebrafish (*Danio rerio*) pathogenicity in *Pseudomonas protegens* CHA0 and *Pseudomonas baetica* LMG25716. **(A)** Kaplan-Meier survival curves of zebrafish embryos (3 days post-fertilization) following bath immersion infection with *Pseudomonas protegens* CHA0, its Fit toxin mutant CHA0 Δ*fitD*, *Pseudomonas baetica* LMG25716 its toxin mutant *P. baetica* Δ*fitD*, or sterile egg water control at a cell density of 10^7^ CFU mL^−1^ over 48 h (n = 10 embryos per condition, three independent biological replicates). Different letters indicate statistically significant differences between treatments (log-rank test). **(B)** Systemic virulence assay following microinjection of 10^3^ CFU into the caudal vein of 2 dpf zebrafish embryos (n = 20 embryos per condition, three independent replicates). Different letters denote statistical significance (log-rank test).

In bath immersion assays performed with 3 days post-fertilization (dpf) embryos, a developmental stage at which the mouth begins to open, *P. protegens* CHA0 caused substantial mortality, reducing survival to less than 40% within 48 h after inoculation (**Figure 8A**). Deletion of *fitD* markedly attenuated this phenotype, with survival of embryos exposed to the Δ*fitD* mutant remaining significantly lower than the wild-type *P. protegens* CHA0. These results indicate that *P. protegens* CHA0 can kill both invertebrate and vertebrate hosts and that FitD contributes to its virulence during waterborne infection of zebrafish embryos.

Exposure to *P. baetica* LMG25716 also caused embryo mortality, with survival decreasing to approximately 60 % after 48 h (**Figure 8A**). However, the Δ*fitD* mutant displayed mortality kinetics similar to the wild-type strain, suggesting that FitD is not required for the virulence of *P. baetica* during waterborne infection under the tested conditions.

Previous work showed that the pathogenicity of *P. baetica* LMG25716 is strongly influenced by the route of infection. In wedge sole (*D. cuneata*), intraperitoneal injection of LMG25716 resulted in 100% mortality, whereas immersion caused only 10 % mortality, consistent with entry through surface wounds or tissue abrasions rather than through ingestion^17^. Based on these published observations, we examined the systemic virulence of *P. baetica* after direct vascular inoculation in zebrafish embryos. We injected the wild-type strains and their Δ*fitD* mutants into the caudal vein of 2 dpf zebrafish embryos (**Figure 8B**). Injection of *P. protegens* CHA0 reduced survival to approximately 75 % at 72 h, whereas survival following injection of the Δ*fitD* mutant was approximately 90 %. This significant attenuation indicates that FitD contributes to systemic virulence in CHA0. *P. baetica* exhibited greater systemic killing activity than CHA0 after direct vascular administration, but no significant difference in survival was observed between wild-type *P. baetica* and its Δ*fitD* mutant (**Figure 8B**).

Together, these results reveal distinct contributions of FitD to zebrafish pathogenicity in the two species. In *P. protegens* CHA0, deletion of *fitD* attenuated mortality after both bath exposure and systemic injection, indicating that the toxin contributes to virulence across different routes of infection. By contrast, deletion of *fitD* did not alter the pathogenicity of *P. baetica* LMG25716 in either assay. The virulence of *P. baetica* in zebrafish embryos therefore appears to depend predominantly on factors other than FitD under the conditions tested.

## Conclusion

In summary, this study reshapes our understanding of the distribution, evolution and functional diversification of the Fit toxin cluster within the *Pseudomonas fluorescens* species complex. The complete *fitABCDEFGH* locus, initially characterized as an insecticidal virulent factor of plant-beneficial pseudomonads, is not restricted to this lifestyle: it is also found in the soil-associated Irchel subgroup and in the fish pathogen *Pseudomonas baetica*, and it can contribute to virulence in a vertebrate host. This challenges the view that the toxin system is tied to biocontrol strains and insect hosts.

Our findings indicate that the history of the locus is not one of simple vertical inheritance. The cluster has a patchy distribution, derives from several acquisitions from donors related to *Photorhabdus* and *Xenorhabdus*, occupies subgroup-specific chromosomal contexts, and has been exchanged between subgroups, as illustrated by the *P. chlororaphis*-type locus of *Pseudomonas* sp. CMR5c and CMR12a. Within this conserved genetic unit, variation is concentrated in the N-terminal effector region of the primary toxin FitD, including the replacement of the Mcf1-type effector by a C80 peptidase domain in *P. baetica*, and in the periplasmic sensory domain of the histidine kinase FitF. The functional contrast between *P. protegens* CHA0 and *P. baetica* is consistent with an influence of these variations, together with the ecological context, on the way the system is deployed: FitD is a major virulence factor of CHA0 in zebrafish but contributes only partially to the virulence of *P. baetica*, which is non-pathogenic in *Pieris brassicae*, likely because of colonisation resistance by gut commensals and a lack of *fitABCDE*activation in the hemolymph.

More generally, the *fit* locus illustrates how a conserved toxin gene cluster can diverge in effector content, regulation and genomic context, providing the raw material for the adaptation of bacterial clades to different hosts and niches. Whether this diversification reflects adaptation to specific hosts or is a consequence of the ecological context in which the toxin is expressed remains an open question that the *fit* system is well suited to address. Finally, because the toxin can act in vertebrates, the host range of *fit*-carrying strains considered for biocontrol should be assessed experimentally rather than inferred from the lifestyle of the producing strain.

## Material and methods

### Bacterial strains, plasmids and culture conditions

All bacterial strains used in this study are listed in **Supplementary Table 1**. All *Pseudomonas* strains were grown in Nutrient Yeast Broth (NYB), incubated overnight at 25 °C with agitation at 180 rpm. *Escherichia coli* strains were grown in Lysogeny broth medium (LB) at 37 °C with shaking (180 rpm). When needed, culture media were supplemented with kanamycin at 25 μg.mL^−1^, gentamycin at 10 μg.mL^−1^.

### Genome dataset and species phylogeny

The presence of the *fit* cluster was assessed by tBLASTn searches of the Fit proteins of *P. protegens* CHA0 (FitABCDEFGH) against all genomes (coverage ≥ 50%, identity ≥ 70%). A genome was considered positive for the Fit cluster only when all eight genes were detected at a single locus and arranged in the conserved gene order. Following this step, genomes of *Pseudomonas* containing the Fit cluster were retrieved from NCBI as well as genomes of representative *Pseudomonas* from the other known *Pseudomonas* subgroup from the *Pseudomonas fluorescens* complex (accession numbers in **Supplementary Table 2**). *Pseudomonas aeruginosa* PAO1 was used as outgroup. A phylogenomic tree was inferred from a concatenated amino acid alignment of single-copy orthologous groups identified using OrthoFinder^28^. The alignment comprised 70 sequences with 464420 amino-acid sites. Maximum-likelihood phylogenetic inference was performed using IQ-TREE v1.6.12^29^. The best-fitting amino acid substitution model, JTTDCMut+F+I+G4, was selected using ModelFinder according to the Bayesian information criterion (BIC). Branch support was assessed using 1,000 ultrafast bootstrap replicates and the SH-like approximate likelihood ratio test (SH-aLRT) with 1,000 replicates. The tree was visualized and annotated with the online tool iTOL^30^.

### Synteny analysis

The Fit cluster and its flanking regions were extracted from the GenBank files of the 17 selected genomes carrying the complete cluster with a window of ± 150 kb. Gene cluster comparisons were performed and visualized with clinker^31^ using default parameters, minimum identity 0.3 for links between genes. Homologous genes were grouped by clinker based on sequence similarity. Regions were aligned on the Fit cluster and displayed in the orientation of the cluster. Average amino-acid identity between clusters was calculated as the average of the pairwise identities reported by clinker between homologous Fit proteins (the eight genes, each counted once and not weighted by length) for each pair of strains and summarized by subgroup.

### Phylogenetic analysis of the Fit toxin-associated gene cluster

Phylogenetic analyses were performed independently for each of the eight proteins encoded by the *fitABCDEFGH* gene cluster. Individual amino acid sequences were aligned using MAFFT^32^. The resulting alignments were curated using ClipKIT^33^ with the smart-gap trimming mode. Maximum-likelihood phylogenetic trees were inferred from the curated alignments using IQ-TREE^29^, with automatic selection of the best-fitting amino acid substitution model and 1,000 ultrafast bootstrap replicates.

To investigate the evolutionary origin of FitD, the corresponding phylogenetic analysis was extended to include Mcf toxin amino acid sequences from *Photorhabdus* and *Xenorhabdus*. Genome accession numbers for the additional sequences are provided in **Supplementary Table 2**. This comparative analysis was used to assess the phylogenetic relationships between FitD and MCF toxins and to explore their possible evolutionary relatedness.

### Protein domain and topology analysis

Domains of FitD and FitF were identified with InterProScan (version 6, ^34^) against the Pfam, SMART, CDD, and PANTHER databases. Identity between the periplasmic sensor domains was calculated from pairwise alignments with EMBOSS Needle online tool^35^.

### Construction of *fitD* deletion mutants and *fit* toxin transcriptional reporters of *P. protegens* CHA0 and *P. baetica* LMG25716

*fitD* deletion mutants of *P. protegens* CHA0 and *P. baetica* LMG25716 were constructed using the pEMG suicide vector and the I-SceI system, following the previously described protocol for *P. protegens*^12^. For the preparation of competent *P. baetica* cells, bacteria were grown at 30 °C for 5 h to reach exponential phase. Transcriptional reporters for *fit* toxin expression were constructed by cloning the 400-bp region upstream of *fitABCDE* of both *P. baetica* LMG25716 and *P. protegens* CHA0 into the pOT1e GFP reporter vector, upstream of the *egfp* gene. The resulting plasmids, pOT1e::P*_fit_*__CHA0_ and pOT1e::P_fit_LMG25716_, were transformed into *P. protegens* CHA0 and *P. baetica* LMG25716*. P. protegens* CHA0 and *P. baetica* LMG25716 carrying the empty pOT1e vector were used as negative controls for GFP expression. The plasmids and primers used are listed in **Supplementary Table 3 and 4 respectively**.

### Expression of FitABCDE promoter

Overnight cultures of the reporter strains were restarted into fresh NYB (1:100) supplemented with gentamycin (10 μg mL^−1^). When cells reached exponential growth phase, i.e., at an optical density (OD_600nm_) of 0.4-0.6, the bacterial cells were washed with Minimal Medium (MM; 0,1% [w/v] NH_4_Cl, 3,49% [w/v] Na_2_HPO_4_ · 2H_2_O, and 2,77% [w/v] 192 KH_2_PO_4_). All strains were inoculated at an OD_600nm_ of 0.05 in 200 μL of grace insect medium (GIM, ^36^) per well in a 96-well black microplate with transparent bottom (Greiner BIO-ONE, ref-655090). Non inoculated wells containing 200 μL of GIM were used to obtain blank values. GFP fluorescence (excitation 479 nm, emission 520 nm) and OD_600nm_ were then monitored every 15 min for 24 h. The data obtained were normalized with the fluorescence and OD_600nm_ blank values and used to calculate the fluorescent signal related to the cell density, *i.e.* Relative Fluorescent Unit (RFU) = (GFP_value_ – GFP_blank_) / (OD_value_ – OD_blank_). The experiment was repeated twice.

### Interbacterial *in vitro* competition assays

The competitive performance of *P. protegens* CHA0 and *P. baetica* LMG25716 was evaluated on solid medium against *Enterobacter* sp. and *Enterococcus* sp., two commensal isolates obtained from the gut of *Pieris brassicae* (^24^; **Supplementary Table 1**). Overnight cultures were centrifuged (5000 *g* for 2 min), washed and resuspended in 0.9% NaCl. Each suspension was adjusted to an OD_600nm_ of 0.1. A *Pseudomonas* strain culture and one gut isolate culture were then combined at a 1:1 ratio, and the mixture was applied to a sterile cellulose acetate membrane filter (0.2 µm pore size, 25 mm diameter; Sartorius) positioned onto nutrient agar medium. Following incubation for 24 h at 25 °C, each filter was transferred to a 15 mL tube containing 5 mL sterile 0.9 % NaCl and thoroughly vortexed to recover the cells. Serial dilutions were subsequently plated under selective conditions to enumerate each competitor. In assays involving *Enterococcus* sp., the commensal was recovered on nutrient agar containing gentamycin (10 µg mL^−1^), whereas CHA0 and LMG25716 were selected on nutrient agar supplemented with ampicillin (100 µg mL^−1^); all plates were incubated at 25 °C. For competitions with *Enterobacter* sp., the commensal was enumerated on antibiotic-free nutrient agar incubated at 42 °C, while the *Pseudomonas* strains were recovered on nutrient agar containing spectinomycin (50 µg mL^−1^) at 25 °C. Colony-forming units of both competitors were determined at 0 and 24 h. The competitive index (CI) was calculated as previously described^24^: CI = (CFU*_Pseudomonas_*_, 24 h_/CFU_gut isolate, 24 h_)/(CFU*_Pseudomonas_*_, 0 h_/CFU_gut isolate, 0 h_). Each competition included four replicate filters, and all assays were repeated in three independent experiments.

### *Pieris brassicae* pathogenicity assays

The pathogenic potential of *P. protegens* CHA0 and *P. baetica* LMG25716 toward *Pieris brassicae* was examined using oral-exposure and hemocoel-injection assays. Newly hatched larvae were reared on pesticide-free cabbage in a Percival PGC-7L2 growth chamber maintained at 25 °C and 60 % relative humidity under a 16 h light (light intensity, 160 μE m^−2^ s^−1^) / 8 h dark photoperiod. For each oral-exposure condition, 18 second-instar larvae (1.0-1.5 cm long) were starved overnight and individually placed in six-well plates. Each larva received a 0.6 g portion of artificial diet containing horseradish powder as a feeding stimulant^12^. The diet was supplemented with 5 μL of washed bacterial suspension containing 5.0 × 10^6^ cells in sterile 0.9% NaCl; control larvae received diet treated with an equal volume of sterile 0.9 % NaCl. Individuals that did not consume the entire portion were omitted from the analysis. After 24 h, larvae were transferred from the plates to Petri dishes, supplied with fresh sterile diet, and scored for survival at 24-h intervals for seven days. Five independent experiments were conducted.

For systemic infection, 2.5 μL of bacterial suspension containing 10^2^ washed cells was delivered into the hemocoel through the second proleg of fourth-instar *P. brassicae* larvae (2.5-3.0 cm long). Each treatment comprised 18 larvae per experiment. Following inoculation, larvae were maintained in the growth chamber in Petri dishes at three individuals per dish. Survival was recorded hourly from 19 h post-injection onward. The assay was performed in three independent experiments.

### *Pieris brassicae* colonization assays

For the *P. brassicae* colonization experiments, *P. protegens* CHA0 and *P. baetica* LMG25716 were chromosomally labelled with a constitutively expressed GFP marker using pBK-miniTn*7-gfp2* as previously described^4^. Larvae were infected orally as described above, with two modifications: third-instar larvae (2.0-2.5 cm in length) were used, and each larva received 10 μL containing 10^7^ bacterial cells. At 24 h post oral infection, larvae were chilled on ice. Hemolymph was discarded after cutting a proleg, and the complete intestinal tract was subsequently collected and transferred into a tube containing 900 μL sterile 0.9% NaCl and sterile glass beads (1 mm, Sigma-Aldrich, G1152). Samples were homogenized with a FastPrep-24 5G instrument (MP Biomedicals) for two 40 s cycles at 6 m s^−1^. Gut homogenates were serially diluted, and 10 μl aliquots were spotted onto nutrient agar supplemented with gentamycin (10 µg ml^−1^). Colony-forming units were quantified using a Fusion FX Spectra imaging system (Vilber-Lourmat®); colonies derived from the inoculated strains were identified by GFP fluorescence under blue-light illumination (∼470 nm). Three independent experiments were performed.

### Zebrafish husbandry, embryo care and infection experiments

Zebrafish were maintained and raised under standard conditions, and according to Swiss regulations (Canton Vaud, license number VD-H28). Adult transparent adult *mitfa^w^*^2^*^/w^*^2^; *mpv17^b^*^18^*^/b^*^18^ (*casper)* mutant zebrafish were maintained at 26 °C in aerated tanks with 10/14 h dark/light cycle ^37^. Zebrafish embryos were collected within the first hours post-fertilization and maintained in E3 medium (5 mM of NaCl; 0.17 mM of KCl; 0.33 mM of CaCl_2_ · H_2_O; 0.33 mM of MgCl_2_ · 7H_2_O, and supplemented with 0.3 mg mL^−1^ methylene blue) and incubated at 28 °C. Before systemic injection or bath infection, embryos were mechanically dechorionated if necessary.

### Bath infection of *Danio rerio* embryos with *Pseudomonas* strains

Overnight cultures of *P. protegens* CHA0, *P. baetica* LMG25716 and their respective mutants were restarted 1:100 in 8 mL of fresh NYB and incubated at 25 °C, 180 rpm for 4 h. Then, exponential growth stage bacteria were centrifuged and resuspended in sterile filtered egg water medium (60 µg mL^−1^ sea salts (Sigma-Aldrich, S9883) in demi water) and the OD_600nm_ was adjusted to obtain the desired bacterial concentrations. To infect the zebrafish embryos (3 days post fertilization, dpf), 200 µL of the respective bacterial suspensions was then added to 1.8 mL egg water containing 10 embryos per condition per well in a 12-well plate and kept at 28 °C. As a control, 200 µL of egg water medium instead of a bacterial suspension was added. To count bacteria present for each condition, 20 µL of the water containing the fish were collected and quantified by serial dilution. The mortality was assessed daily by monitoring live and dead embryos at 24 and 48 hours post-infection. This experiment was repeated 3 times independently.

### Systemic infection of *Danio rerio* embryos by caudal vein injection

Bacterial suspensions were prepared as detailed above. Prior to injection, bacteria were harvested by centrifugation and after removal of the supernatant, the pelleted cells were resuspended in 0.25 % (w/v) amaranth (Sigma-Aldrich, A1016) in PBS to aid visualization of the injection process. Zebrafish embryos were then anesthetized in 0.02% (w/v) buffered 3-aminobenzoic acid methyl ester (pH 7.0) (Tricaine; Sigma-Aldrich, A5040). One nanoliter of the bacterial suspension containing approximately 10^2^ or 10^3^ CFU was individually injected by microinjection into the caudal vein of *casper* zebrafish embryos at 2 dpf. To determine the exact number of bacteria injected, quantitative plating of the injection volume was performed on NA plates supplemented with gentamycin (10 µg mL^−1^). In total 20 zebrafish were injected per condition and kept in 4 mL of egg water in a 6-well plate at 28 °C. The mortality of the embryos was assessed daily and determined visually based on the absence of heartbeats at 24, 48 and 72 hours post-infection. This experiment was repeated 4 times independently.

### Statistical analysis

All the data were analyzed using R studio (version 2026.09.0+174) running R version 4.6.1 and considered significant when *P* < 0.05.

Bacterial load in the insect gut was expressed as CFU per mg of gut and log₁₀-transformed. Samples with no detectable colonies (ND, below the detection limit) cannot be log-transformed and were therefore excluded from the quantitative comparisons, while being kept in the sample counts. Data from three independent experiments were pooled. The log₁₀ CFU values of the two strains among detected samples were compared with a two-sided Mann-Whitney U test (Welch’s t-test gave consistent results). The proportion of colonized insects (detected vs. ND) was compared between strains with a two-sided Fisher’s exact test and represented as stacked bar plots.

Competitive ability of *P. baetica* LMG25716 and *P. protegens* CHA0 was assessed against *Enterobacter* sp. and *Enterococcus* sp. Competitive indices of LMG25716 and CHA0 were compared against each competitor with a two-sided Welch’s t-test.

Survival after oral exposure (insects) and after injection (insects, zebrafish), and after bath exposure (zebrafish), was analyzed with Kaplan–Meier estimators, individuals alive at the end of follow-up being right-censored. Independent experiments were pooled within each assay (insect feeding: 5 experiments; insect injection: 3 experiments; zebrafish injection: 4 experiments; zebrafish bath exposure: 3 experiments), and each assay was analyzed separately. Survival curves of all treatments were first compared with a global log-rank test, then with pairwise log-rank tests corrected for multiple comparisons by the Holm method (α = 0.05). In insect assays, each treatment was compared with the NaCl control; in zebrafish assays, all pairs of treatments were tested. Hazard ratios (HR) and 95 % confidence intervals were estimated with Cox proportional hazards models stratified by experiment, with a ridge penalty (0.01) to stabilize estimates in groups with few or no events; groups without any death (controls) were excluded from the Cox models because their HR is not estimable and were compared by log-rank test only. In zebrafish experiments, CHA0 was used as the reference in Cox models. Heterogeneity between experiments was assessed with a likelihood-ratio test of the treatment × experiment interaction, and robustness was evaluated by repeating the pairwise comparisons after removing each experiment in turn. For the zebrafish bath assay, where follow-up was 48 h for all fish, final mortality was also compared between treatments with two-sided Fisher’s exact tests corrected by the Holm method.

## Acknowledgments

We are grateful to Dr. José A. López for kindly providing the *Pseudomonas baetica* type strain LMG25716. We also thank Prof. Rolf Kümmerli (University of Zurich) for generously sharing the environmental Irchel *Pseudomonas* strains. We would like to thank Samantha Gozel and Beatrice Tappi for their technical assistance. Finally, we express our warm gratitude to the laboratory support staff and media preparation facility (Nazife Beqa, Latifa Labidi, Nelson Galvao) for their outstanding technical assistance in preparing culture media.

The first author dedicates this work to the memory of her beloved sister Virginie, who now shines among the stars – 14/11/2026.

## Data availability

Dataset and raw data were deposited on Zenodo (10.5281/zenodo.23888124).

## Sources of support (grants/ equipment)

This study was supported by the Swiss National Science Foundation (SNSF) (grant no. 310030_184666), by the SNSF-funded Swiss National Centre of Competence in Research (NCCR) Microbiomes (grant no. 19 51NF40_180575 and 51NF40_225148), and the University of Lausanne, Switzerland.

