## Supplementary tables and figures for "Modular evolution and functional divergence of the *Pseudomonas* Fit toxin locus"

#### Content:

##### Supplementary Figures

**Supplementary Figure 1:** *Single-gene phylogenetic trees of individual components of the fit locus (FitA – FitG) across Pseudomonas subgroups.*

**Supplementary Figure 2:** *Oral and systemic toxicity of Pseudomonas baetica LMG25716 and environmental Irchel pseudomonads towards Pieris brassicae larvae.*

##### Supplementary Tables

**Supplementary Table 1:** *Bacterial strains used in this study.*

**Supplementary Table 2:** *List of all genomes used in this study.*

**Supplementary Table 3**: *Plasmids used in this study.*

**Supplementary Table 4**: *Primers used in this study.*

**References**

### Supplementary Figures

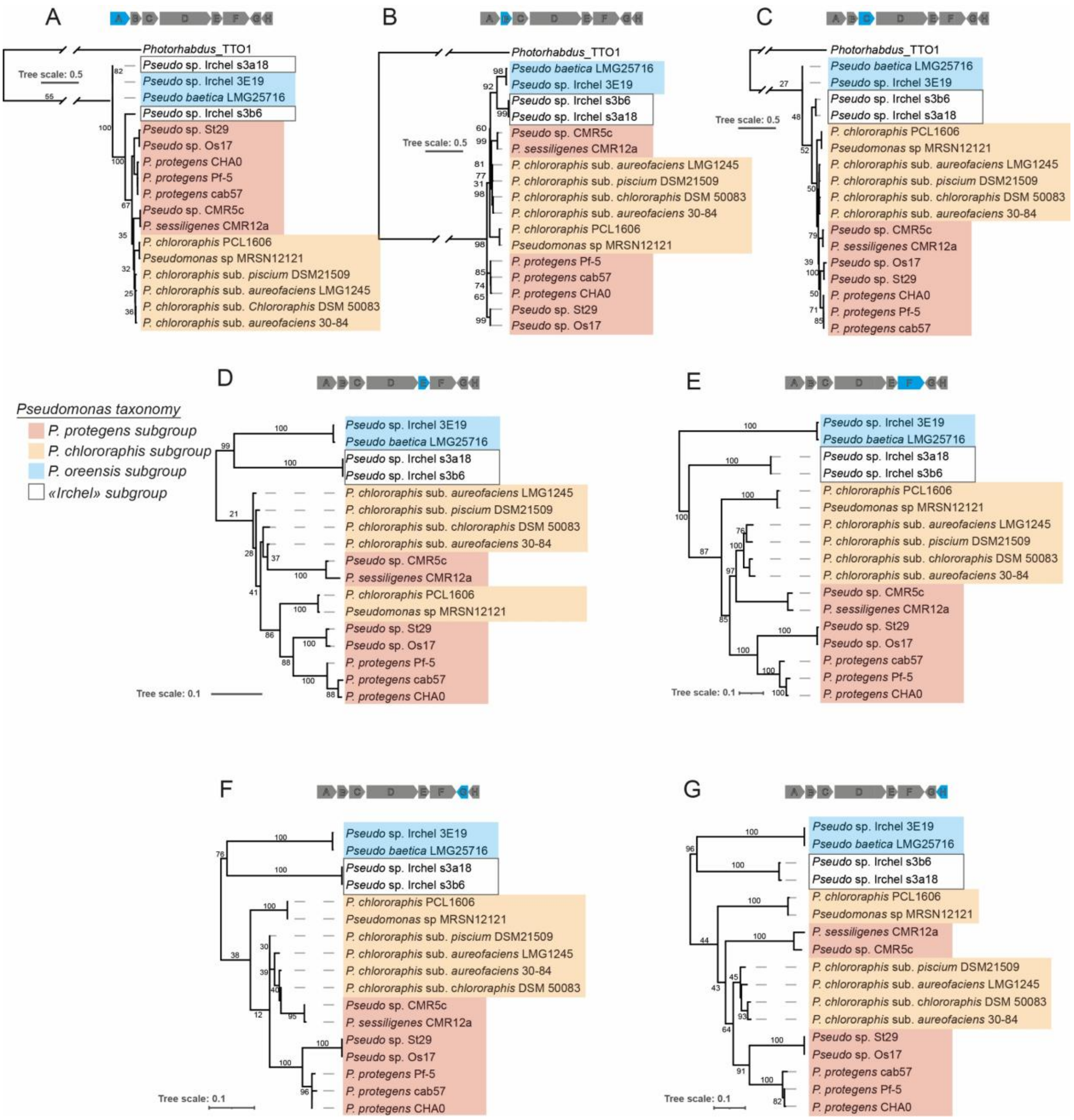


**Supplementary Figure 1: Single-gene phylogenetic trees of individual components of the fit locus (FitA – FitG) across *Pseudomonas* subgroups.** Maximum-likelihood phylogenetic trees based on the alignment of amino acid sequences of **(A)** FitA, **(B)** FitB, **(C)** FitC, **(D)** FitE, **(E)** FitF, **(F)** FitG, and **(G)** FitH.

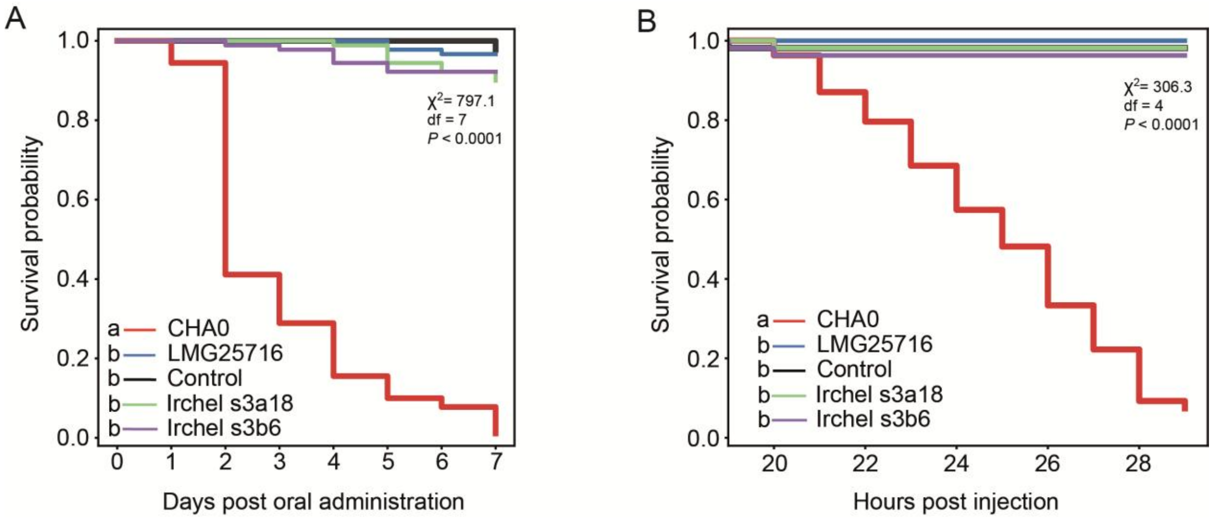


**Supplementary Figure 2: Oral and systemic toxicity of *Pseudomonas baetica* LMG25716 and environmental Irchel pseudomonads towards *Pieris brassicae* larvae. (A)** Kaplan-Meier survival curves of *Pieris brassicae* following oral administration of *P. protegens* CHA0, *P. baetica* LMG25716, *Pseudomonas* sp. Irchel s3a18, *Pseudomonas* sp. Irchel s3b6 or control. Statistical significance was assessed using a log-rank test, different letters indicate statistically significant differences at $P < 0.05$. **(B)** Kaplan-Meier survival curves of *P. brassicae* larvae following direct microinjection into the hemolymph of the same bacterial strains.

### Supplementary Tables

**Supplementary Table 1:** Bacterial strains used in this study.

| **Strains** [1] | **Description** [2] | **Reference or source** |
|---|---|---|
| ***Pseudomonas strains*** | | |
| *P. protegens* CHA0$^T$ | Wild-type, isolated from tabaco roots | 1 |
| *P. baetica* LMG25716$^T$ | Wild-type, isolated from edge sole *Dicologlossa cuneata* | 2 |
| *Pseudomonas* sp. Irchel s3a18 | Wild type, isolated from soil | 3 |
| *Pseudomonas* sp. Irchel s3b6 | Wild type, isolated from soil | 3 |
| ***Pseudomonas derivatives*** | | |
| LMG25716 Δ*fitD* | *fitD* in frame deletion mutant of CHA0 (ATI02_1887) | This study |
| LMG25716 pOT1e::$P_{fit\_LMG25716}$ | LMG25716 harboring the transcriptional reporter vector pOT1e::$P_{fit\_LMG25716}$, $Gm^R$ | This study |
| CHA0 Δ*fitD* | In frame deletion mutant of *fitD* (PPRCHA0_3005) | This study |
| CHA0 pOT1e::$P_{fit\_CHA0}$ | CHA0 harboring the transcriptional reporter vector pOT1e::$P_{fit\_CHA0}$, $Gm^R$ | This study |
| CHA0-*gfp2* | CHA0::attTn*7*-*gfp2*; $Gm^r$ | 4 |
| LMG25716-*gfp2* | LMG25716::attTn*7*-*gfp2*; $Gm^r$ | This study |
| ***Insect gut isolates*** | | |
| *Enterobacter* sp. | Isolated from the gut of *Pieris brassicae* | 5 |
| *Enterococcus* sp. | Isolated from the gut of *Pieris brassicae* | 5 |
| ***Escherichia coli*** | | |
| *E. coli S17-1/λpir* | Laboratory strain | 6 |
| *E. coli DH5α* | Laboratory strain | 7 |

[1] $^T$, type strain
[2] : $Gm^R$ : Gentamycin resistant

**Supplementary Table 2:** List of all genomes used in this study.

| **Genome accession numbers** | **Strain name** |
|---|---|
| | ***Photorhabdus* spp.** |
| GCA_000196475.1 | *Photorhabdus asymbiotica* ATCC43949 |
| GCA_000711895.1 | *Photorhabdus australis* DSM 17609 |
| GCF_000196155.1 | *Photorhabdus laumondii* TTO1 |
| GCA_000722995.1 | *Photorhabdus temperata* subsp. *temperata* Meg1 |
| | ***Providencia* spp.** |
| GCF_002393505.1 | *Providencia alcalifaciens* FDAARGOS_408 |
| | ***Pseudomonas* spp.** |
| GCF_001045685.1 | *Pseudomonas aeruginosa* DSM 50071 |
| GCF_900109755.1 | *Pseudomonas agarici* LMG 2112 |
| GCF_900105475.1 | *Pseudomonas asplenii* ATCC 23835 |
| GCF_001983205.1 | *Pseudomonas azotoformans* DSM 18862 |
| GCF_019733355.1 | *Pseudomonas baetica* 25P2F9 |
| GCA_031583675.1 | *Pseudomonas baetica* K12 |

| | |
|---|---|
| GCF_002813455.1 | *Pseudomonas baetica* LMG 25716 |
| GCF_012034345.1 | *Pseudomonas brassicacearum* JCM 11938 |
| GCF_900105815.1 | *Pseudomonas brenneri* DSM 15294 |
| GCF_000963835.2 | *Pseudomonas chlororaphis* PCL1606 |
| GCF_003851835.1 | *Pseudomonas chlororaphis* subsp. *aurantiaca DSM 19603* |
| GCF_000761195.1 | *Pseudomonas chlororaphis* subsp. *aurantiaca* JD37 |
| GCF_003851165.1 | *Pseudomonas chlororaphis* subsp. *aurantiaca* M12 |
| GCF_002355875.1 | *Pseudomonas chlororaphis* subsp. *aurantiaca* StFRB5008 |
| GCF_000281915.1 | *Pseudomonas chlororaphis* subsp. *aureofaciens* 30-84 |
| GCA_001269595.1 | *Pseudomonas chlororaphis* subsp. *aureofaciens* CD |
| GCF_003851905.1 | *Pseudomonas chlororaphis* subsp. *aureofaciens* DSM 6698 |
| GCF_000698865.1 | *Pseudomonas chlororaphis* subsp. *aureofaciens* PA23 |
| GCF_009834495.1 | *Pseudomonas chlororaphis* subsp. *aureofaciens* R47 |
| GCF_001602135.1 | *Pseudomonas chlororaphis* subsp. *chlororaphis* 189 |
| GCA_003851145.1 | *Pseudomonas chlororaphis* subsp. *chlororaphis* ATCC17415 |
| GCF_003945765.1 | *Pseudomonas chlororaphis* subsp. *chlororaphis* DSM 50083 |
| GCF_900115645.1 | *Pseudomonas chlororaphis* subsp. *chlororaphis* NFPP07 |
| GCF_900104995.1 | *Pseudomonas chlororaphis* subsp. *piscium* DSM 21509 |
| GCF_000597925.1 | *Pseudomonas chlororaphis* subsp. *piscium* HT66 |
| GCF_003850445.1 | *Pseudomonas chlororaphis* subsp. *piscium* PCL1391 |
| GCF_900106085.1 | *Pseudomonas corrugata* LMG 2172 |
| GCF_900215245.1 | *Pseudomonas fluorescens* ATCC 13525 |
| GCF_900105835.1 | *Pseudomonas fragi* NRRL B-727 |
| GCF_900105495.1 | *Pseudomonas frederiksbergensis* LMG 19851 |
| GCF_001983165.1 | *Pseudomonas gessardii* DSM 17152 |
| GCF_900105485.1 | *Pseudomonas granadensis* LMG 27940 |
| GCF_900182985.1 | *Pseudomonas helmanticensis* LMG 28168 |
| GCF_002236115.1 | *Pseudomonas jessenii* DSM 17150 |
| GCF_900105635.1 | *Pseudomonas kilonensis* DSM 13647 |
| GCF_024169245.1 | *Pseudomonas koreensis* Ps 9-14 |
| GCF_900104735.1 | *Pseudomonas lini* DSM 16768 |
| GCF_900103705.1 | *Pseudomonas lundensis* LMG 13517 |
| GCF_002091695.1 | *Pseudomonas mandelii* NBRC 103147 |
| GCF_900105325.1 | *Pseudomonas marginalis* NCPPB 667 |
| GCF_900106005.1 | *Pseudomonas mediterranea* DSM 16733 |
| GCF_024169545.1 | *Pseudomonas migulae* CFML 95-321 |
| GCF_900102045.1 | *Pseudomonas moorei* DSM 12647 |
| GCF_900106045.1 | *Pseudomonas mucidolens* LMG 2223 |
| GCF_900105795.1 | *Pseudomonas orientalis* LMG 23660 |
| GCF_900105155.1 | *Pseudomonas prosekii* LMG 26867 |
| GCF_000828695.1 | *Pseudomonas protegens* Cab57 |
| GCF_900560965.1 | *Pseudomonas protegens* CHA0 |
| GCF_000012265.1 | *Pseudomonas protegens* Pf-5 |
| GCF_007858275.1 | *Pseudomonas proteolytica* DSM 15321 |
| GCF_001043005.1 | *Pseudomonas psychrophila* DSM 17535 |
| GCF_900105185.1 | *Pseudomonas saponiphila* DSM 9751 |

| | |
|---|---|
| GCF_003850565.1 | *Pseudomonas sessilinigenes* CMR12a |
| GCF_900111895.1 | *Pseudomonas simiae* CCUG 50988 |
| GCF_003850545.1 | *Pseudomonas* sp. CMR5c |
| GCF_900187585.1 | *Pseudomonas* sp. Irchel 3E19 |
| GCF_900187535.1 | *Pseudomonas* sp. Irchels3a18 |
| GCF_900187415.1 | *Pseudomonas* sp. Irchels3b6 |
| GCF_009861525.1 | *Pseudomonas* sp. LD120 |
| GCF_000931465.1 | *Pseudomonas* sp. MRSN12121 |
| GCF_001547895.1 | *Pseudomonas* sp. Os17 |
| GCF_001547915.1 | *Pseudomonas* sp. St29 |
| GCF_900105675.1 | *Pseudomonas synxantha* LMG 2190 |
| GCF_900104825.1 | *Pseudomonas taetrolens* LMG 2336 |
| GCF_900102295.1 | *Pseudomonas thivervalensis* DSM 13194 |
| GCF_002813445.1 | *Pseudomonas tolaasii* NCPPB 2192 |
| GCF_900105285.1 | *Pseudomonas umsongensis* LMG 21317 |
| GCF_008801375.1 | *Pseudomonas vancouverensis* CCUG 49675 |
| GCF_001043055.1 | *Pseudomonas weihenstephanensis* DSM 29166 |
| GCF_900105735.1 | *Pseudomonas yamanorum* LMG 27247 |
| **_Xenorhabdus_ spp.** | |
| GCF_002632465.1 | *Xenorhabdus budapestensis* DSM 16342 |
| GCF_002632725.1 | *Xenorhabdus hominickii* DSM 17903 |
| GCF_900113945.1 | *Xenorhabdus mauleonii* DSM 17908 |
| GCA_002632615.1 | *Xenorhabdus miraniensis* DSM 17902 |
| GCF_000953355.1 | *Xenorhabdus nematophila* AN6/1 |
| GCA_002632825.1 | *Xenorhabdus stockiae* DSM 17904 |
| GCF_002127535.1 | *Xenorhabdus vietnamensis* DSM 22392 |

**Supplementary Table 3:** Plasmids used in this study.

| **Plasmid** | **Genotype or relevant characteristics**[1] | **Reference or source** |
|---|---|---|
| pEMG | Expression vector; oriR6K, lacZα with two flanking I-SceI sites; $Km^R$, $Ap^R$ | [4] |
| pOT1e | Promoter-probe vector derived from pBBR1; contains promoter-less *egfp*; $Gm^R$ | [8] |
| pBK-miniTn7-gfp2 | pUC19-based delivery plasmid for miniTn7-gfp2; mob+; $Gm^R$, $Cm^R$, $Ap^R$ | [9] |
| pMUT0001 | pEMG::Δ*fitD*; suicide plasmid for the in-frame deletion of the *fitD* gene in *P. baetica* LMG25716 | This study |
| pOT1e::$Pfit_{LMG25716}$ | pOT1e containing *egfp* gene under the control the promoter of the *fit* gene cluster of *P. baetica* LMG25716; $Gm^R$ | |
| pSW-2 | oriRK2, xylS, Pm::I-sceI; $Gm^R$ | [10] |
| pUX-BF13 | Helper plasmid encoding Tn7 transposition functions; R6K-replicon; $Ap^R$ | [11] |

[1] $Ap^R$, ampicillin resistance; $Cm^R$, chloramphenicol resistance; $Gm^R$, gentamycin resistance; $Km^R$, kanamycin resistance; $Tc^R$, tetracycline resistance.

**Supplementary Table 4**: Primers used in this study.

| Primer names | Sequence 5' → 3', [1] | Usage | References |
|---|---|---|---|
| baet_del-fitD-1 | cg**GAATTC**TTTGTCGAGGCGCAAAAAG | Deletion of *fitD* in *Pseudomonas baetica* LMG25716 | This study |
| baet_del-fitD-2 | cg**GGATCC**CATCGCAGGGCGCTCGAT | | |
| baet_del-fitD-3 | cg**GGATCC**CATGGTATGTTGGCCTGTG | | |
| baet_del-fitD-4 | acgc**GTCGAC**TGAGGTAATGAACCCGGGA | | |
| Fitclust-I-F | AAGGTTATCGCGGCTATCT | External primer to control *fitD* removal | This study |
| Fitclust-I-R | CCTGGATGAGCTATGAGA | | |
| Baet_exp-fit-1 | g**GAATTC**CGCAGGGCGCTCGATGGC | Construction of a transcriptional reporter of Fit toxin expression in *P. baetica* | This study |
| Baet_exp-fit-2 | acgc**GTCGAC**CTATACGCCCCGCCTTTA | | |
| pEMG-F | GTTTTCCCAGTCACGACG | pEMG verification of fragment insertion | 12 |
| pEMG-R | GCTTCCGGCTCGTATGTTGT | | |

[1]: Restriction enzymes are indicated in bold and underlined.
